# Growing despite ectoparasite infection: sex-specific effects on telomere shortening in a free-living corvid

**DOI:** 10.64898/2026.10.21.760650

**Authors:** Elisa P. Badás, Christina Bauch, Jelle J. Boonekamp, Ellis Mulder, Simon Verhulst

**Affiliations:** Department of Evolutionary Ecology, National Museum of Natural Sciences - Spanish National Research Council (CSIC). C/ José Gutiérrez Abascal 2, 28006 Madrid, Spain; Groningen Institute for Evolutionary Life Sciences, University of Groningen, Nijenborgh 7, 9747 AG Groningen, Netherlands; School of Biodiversity, One Health & Veterinary Medicine, University of Glasgow, Glasgow, United Kingdom

**Keywords:** cross-foster, environmental effects, host-parasite interactions, longitudinal, TRF

## Abstract

Ectoparasite infections during development are known to impact fitness-related traits, but their physiological costs in relation to sex or growth trade-offs are understudied. In birds, previous studies at the brood level have found costs of infection by blood-sucking ectoparasites in molecular biomarkers (i.e., increased telomere shortening). However, infection probability also varies with age, sex, or within broods. Whether such costs are age or sex-dependent is not well known, and disentangling these effects is of interest, because telomeres potentially reveal the long-term impact of infection. Here, we used 6-year-data from jackdaw nestlings (*Coloeus monedula*) to investigate individual level associations of infection by the fly (*Carnus hemapterus*) at two intermediate time points during development, with telomere dynamics over the nestling period. We evaluated (i), *Carnus* presence in relation to sex, brood size manipulation, body mass, and size, and (ii), how telomere shortening varied in relation to parasite presence and achieved growth. Early in development (day 10), flies were more prevalent on heavier nestlings, while later in development (day 20), they were more prevalent on smaller and lighter nestlings, suggesting different ectoparasite preference. Telomere loss was higher in female nestlings when exposed to parasites only on day 20, and in males, when they were heavy earlier in development. In conclusion, infection is a dynamic process, and ectoparasite effects may depend on the timing of infection and reveal sex-specific growth vs. self-maintenance trade-offs. Our study indicates a need to sample at multiple developmental stages when assessing host-parasite interactions and the physiological costs of infection.

## INTRODUCTION

In vertebrates, telomeres –the TTAGGG_n_ DNA sequence that caps the ends of eukaryotic chromosomes in a repetitive, non-coding manner– contribute to provide DNA stability and integrity with each cell replication (Blackburn 1991). At the cellular level, telomere length is known to be influenced by molecular and physiological processes; for example, cell replication shortens telomeres (Blackburn 2000) and oxidative stress accelerates their shortening (von Zglinicki 2002). These processes partly explain the variability in telomere length across individuals at the organismal level, in both model and non-model species, and such variation is determined not only by genetic (Broer et al. 2013; Bauch et al. 2022) but also by environmental factors (Tobler et al. 2022). Because external influences leave a detectable ‘footprint’ of past experiences in telomeres, in animal ecology, telomere length has become a well-established marker for fitness prospects, including lifespan, reproductive output (Bauch et al. 2013; Eastwood et al. 2019) and survival (Wilbourn et al. 2018). In this context, telomeres have been suggested to act as markers of somatic redundancy, with the repetitive elements accumulating damage until their depletion prompts functional failure (Boonekamp et al. 2013). Therefore, increased telomere shortening early in life may induce cryptic sublethal effects that do not become apparent until later in life when telomeres become critically short (e.g. (Eastwood et al. 2022)).

Disentangling the effects of early-life adversity on adult performance is complicated by interactions between challenges. For example, in avian species, fitness has been separately related to parasitic infections (Dunn et al. 2021) and brood size-dependent nestling competition (Winder et al. 2025); but they may also interact to impact fitness (Moller 1990). Mechanisms underlying the impact of early-life adversity on adult performance are only partly resolved, and telomere dynamics may be one such mechanism. Indeed, a recent meta-analysis in non-model vertebrates showed a strong effect of endoparasite infections on telomere loss (Chatelain et al. 2019). In contrast, whether ectoparasite infection impacts telomere dynamics is less well known (but see (Dudek et al. 2021; Tschirren et al. 2021; Badás et al. 2023; Wolf et al. 2023). However, a relationship between telomere shortening and infestation with ectoparasites is likely in birds and reptiles, since (i) most ectoparasites infecting birds in high numbers suck blood (Badás 2025), inducing accelerated turnover of the nucleated erythrocytes, and (ii), telomeres shorten during cell-division. Blood-sucking parasites could thus, directly, and possibly also indirectly, through other effects on phenotypic state, accelerate telomere shortening, as seen in birds parasitised by blowfly larvae (O’Brien and Dawson 2009; Sun et al. 2020).

Parasitic infections vary, among other things, in transmission routes, prepatent periods and temporal manifestation patterns. For example, vector-borne avian malaria-like infections require a prepatent period to reach the bloodstream during which they are only detectable by molecular methods and not under microscopic examination (Martínez et al. 2018). As a result of these temporal patterns, the effects of malarial parasites on early-life telomere dynamics might be limited to the latest period of development. Ectoparasite infection patterns throughout nestling development may change due to an increase/decrease in parasite numbers with environmental conditions (García-del Río et al. 2025) and/or parental behaviour (e.g. nest sanitation, (Tomás et al. 2012; Cantarero et al. 2013a), but the effect of such changes in infection on early-life telomere shortening is currently unknown. Studies that consider parasite dynamics early in life are needed to increase the understanding of the effects of host-parasite interactions over time.

In addition to dynamic changes in parasite prevalence during development, the rate of telomere shortening may also vary with age, as seen for example in humans, in which shortening rate is higher earlier in development (e.g. Lai et al., 2023). Parasitic infections could thus differentially affect telomere loss over development, depending on the timing of infection during the nestling period. In wild jackdaws, we previously showed that the presence of the parasitic carnid fly *Carnus hemapterus* in the brood at any point during development was related to accelerated telomere shortening when brood size was enlarged, but not when brood size was reduced (Badás et al 2023). *Carnus hemapterus* is a nestling-specific hematophagous fly that can be found parasitizing several medium-sized bird species (Roulin et al. 2003; Hoi et al. 2018), but its presence may vary with nestling age (Roulin et al. 2003). Thus, parasitic infection effects may depend on the age at which infection takes place, and we here test this hypothesis. Focusing on broods that were observed to be infected at any time point during the nestling period, we first investigated how within- and between-brood variation in *Carnus* presence during early and late development was related to brood manipulation, mass, size and sex of the nestlings. We then tested to what extent individual variation in telomere shortening across the infected individuals can be related to *Carnus* presence and growth traits throughout development.

## MATERIALS AND METHODS

### Study site and sampling

We studied a population of wild jackdaws (*Corvus monedul*a) breeding in nest boxes located south of Groningen, the Netherlands (53° 14’N, 6°64’E). These nestboxes have been routinely monitored since 2005 to record information on reproduction and identify both pair-members by their unique colour ring combination. We here use data collected in 2012-2017; in these years *Carnus hemapterus* presence was recorded and brood size was manipulated. The brood size manipulation protocol is described in detail in (Boonekamp et al. 2020); briefly, when the oldest nestling was 4 days old (hatching date = day 0, hereafter, day 5), dyads of broods were matched by clutch size and laying date (±1 day) to create enlarged and reduced broods. Nestlings were cross fostered (and thus moved from their incubation nest to their foster nest) with the aim to create a net manipulation of 2 nestlings up or down, with both manipulated broods containing nestlings hatched in both broods in the dyad (Bauch et al. 2022). Weight (to the nearest 0.1 g) was measured on days 5, 10, 20 and 30, and wing length on days 10, 20 and 30. On days 5 and 30, a blood sample was taken from the brachial vein. All blood samples were stored in 2% EDTA buffer at 4– 7°C and snap frozen in a 40% glycerol buffer for long-term storage at −80°C within 3 weeks.

Presence of *Carnus hemapterus* (hereafter, *Carnus*) was routinely recorded for each nestling in the brood for each visit. However, *Carnus* were rarely encountered on day 5 and never on day 30, and we therefore restrict our analyses to ectoparasite presence on days 10 and 20 (note that previous studies have reported that carnid flies are likely to move between nestlings within the brood throughout development, Roulin et al., 2003). *Carnus* parasites were either located by visual inspection of the nestlings’ body, or by the identification of bite marks (under the wings, and legs and belly). We previously validated the bite score as a reliable indicator of parasite presence in the nest (Badás et al. 2023), and we therefore assumed that the focal nestling was parasitized when either the *Carnus* parasite count or the bite score index was >0. With the exception of the occasional louse fly *Hippoboscidae*, we did not encounter other ectoparasites, but note that we did not apply dedicated techniques to survey all possible ectoparasites.

#### Telomere length

Telomere length of nucleated erythrocytes was measured using the Telomere Restriction Fragment (TRF) analysis as previously described (Salomons et al. 2009). Briefly, DNA was extracted (CHEF Genomic DNA Plug kit, Bio-Rad, Hercules, CA), digested overnight with proteinase K at 50°C, and about half of the digested DNA was simultaneously digested with Hind III (60U), Hinf I (30U) and Msp I (60U) for 18 h at 37°C in NEB2 buffer. The DNA was then separated by pulsed-field gel electrophoresis at 14°C for 24 h (3 V / cm, initial switch time 0.5 s, final switch time 7.0 s), and gels were dried (Bio-Rad model 538) and hybridized overnight using a ^32^P-end-labelled oligo (5’-CCCTAA-3’)_4_ that binds to the 3’ end-cap telomere overhang. Subsequently, we detected the radioactive signal obtaining a gel picture with a phosphor screen (Cyclone Storage Phosphor System, PerkinElmer) and using IMAGEJ v. 1.38x we quantified the individual telomere length distribution for each sample through densitometry (Bauch et al. 2022). As in Badás et al (2023), for each individual, we used the averaged value of the telomere length size distributions. Same-brood nestlings were spread over gels, and samples from the same individual (days 5 and 30) were always placed in neighbouring lanes (total N = 184 blood samples distributed over 34 gels). The intergel repeatability was measured as the coefficient of variation for a 29-day-old jackdaw control sample that was included in all 34 gels (6.01%), while the intragel repeatability is demonstrated by the high repeatability within-individuals (see Badás et al 2023).

#### Statistical analyses

Statistical analyses were performed in R 4.4.2 (R Development Core Team 2024). We evaluated parasite presence in relation to nestling age, brood size manipulation, body mass, size and sex before testing for parasitic infection effects on telomere dynamics. Because we here focus on within brood variation, we only included individuals from the infected broods in this analysis (but see Tables S1 and S2 for the models including all individuals from the infected and non-infected broods).

To analyse variation in *Carnus* presence/absence throughout development, we used two generalised mixed effects models (GLMM) with binomial error distribution in which *Carnus* presence/absence on days 10 and 20 were the dependent variables (hereafter, D10 and D20 *Carnus* presence models, n = 763 and n = 700 nestlings respectively). For each, we present the model that incorporated all predictors relevant to our hypotheses. The following terms were included as fixed effects: body mass or wing length (measured on days 10 and 20), brood size manipulation category (enlarged vs. reduced), sex, the 2-way interaction among each of these, and brood size on day 5 (when the manipulation took place, see above). Brood size manipulation category and sex were mean centred, to aid in the interpretation of interaction terms (Schielzeth 2010), and continuous variables were standardised (mean = 0, SD = 1). Nestling state (mass, size) varied between and within broods, and these levels of variation may affect *Carnus* presence differently. Therefore, the models included: mean mass and mean wing length in the brood (between-brood effect, hereafter μ mass and μ wing length)and the individual deviation from the brood mean (within-brood effect, hereafter δ mass and δ wing length) as separate variables. Standardization of μ mass and μ wing length was done separately for nestlings in reduced and enlarged broods to avoid confounding brood size manipulation with nestling mass or size (Badás et al 2023).

Day 20 mass correlated strongly with wing length, and thus including both mass and wing length in the D20 *Carnus* presence model resulted in high variance inflation factors (VIF > 5). Preliminary analyses revealed wing length to be a better predictor of *Carnus* presence than mass, and in the D20 model we therefore use wing length instead of mass (there are no day 10 wing length data as the primaries are barely visible at this age). However, for comparison, in Table S3 we show the analogous model to the D10 *Carnus* presence model including D20 mass instead of wing length.

Random effects in the D10 and D20 *Carnus* presence models included: year of birth to control for temporal variation, colony to control for spatial variation, and foster and incubation nest. GLMMs were fitted using the R package *blme* 1.0-4 (Chung et al. 2013) to avoid convergence and singular fit problems. Models fitted with *blme* use a Bayes modal estimation with an inverse Wishart covariance prior for the random effects (Chung et al. 2013). All diagnostics plots were examined to confirm that there were no deviations from model assumptions before interpreting model estimates and all VIF < 5 confirming that there was no collinearity between independent variables (Zuur et al. 2010). In the model tables (results section), we report parameter estimates as the logarithmic of odds ratio. However, in the main text, for ease of interpretation, and because odds ratio do not exactly correspond to prevalence risk ratios or probability (Davies et al. 1998), we report the marginal effects extracted from the R package *ggeffects* 2.2.0 (Lüdecke 2018).

To explore variation in telomere shortening with parasite infection dynamics during development, we filtered the telomere dataset to include the infected broods only. Data on *Carnus* presence/absence per nestling on days 10 and 20 and telomere shortening was available for a subset (n = 186 nestlings) from our previous study (Badás et al 2023), in which we showed synergistic effects between the brood manipulation and *Carnus* infection. We built a linear mixed effects model using the R package *lme4* 1.1-36 (Bates et al. 2014) with telomere shortening as dependent variable (hereafter, telomere model). Telomere shortening covered the entire nestling development because it was calculated as the difference between telomere length at days 5 (shortly after hatching) and 30 (shortly before fledgling). As fixed effects, the telomere model specified: brood manipulation, sex, *Carnus* presence on D10, *Carnus* presence on D20, and the 3-way interactions: *Carnus* presence on D10 x brood manipulation x sex, and *Carnus* presence on D20 x brood manipulation x sex. We specified the same random effects as in the D10 and D20 *Carnus* presence models, with the addition of telomere gel identity to control for between-gel differences. Between-group contrasts in this telomere model were evaluated using the R package *emmeans* 2.0-3 (Lenth and Piaskowski 2026).

We additionally tested whether growth traits explain variation in telomere shortening by building a model of telomere shortening that included δ mass on D10 and δ wing length on D20, with the addition of the same fixed effects and 3-way interactions as described above. Based on our findings (see below), δ mass on D10 and δ wing length on D20 served as proxies for *Carnus* presence early and later during development. None of these telomere models incurred in high VIF. The two telomere models address two biologically different questions (parasite exposure *vs.* growth effects on telomere dynamics) and are therefore not subject to multiple-testing, but for each, we show the corrected AIC (Akaike Information Criterion) and BIC (Bayesian Information Criterion) extracted using the R package *MuMIn* 1.48-19 (Bartoń 2026).

## RESULTS

### Parasite presence during development

Carnid flies were recorded on 54% of 10-day-old nestlings (n = 763). The likelihood of *Carnus* infection increased significantly with increasing nestling mass (Table 1, Figs. 1a & b). Effect estimates of within- and between brood variation in nestling mass were indistinguishable (Table 1), indicating absolute mass predicted the likelihood of being parasitized, independent of the mass of other nestlings in the brood. However, the within-brood mass effect tended to be sex dependent, with *Carnus* parasites found more often on female nestlings above average mass (Fig. 1a): the predicted probability of infection in female nestlings increased from 19% [lower CI = 13%, upper CI = 25%] on chicks weighing one standard deviation (SD=17.32 g) less than the brood mean to 74% [63%, 83%] on female nestlings weighing 1 SD more than the brood mean, and the corresponding numbers for male nestlings were 17% [12%, 22%] and 43% [28%, 59%]. *Carnus* prevalence at the mean weight within-broods was 45% [30%, 61%] for females and 28% [19%, 40%] for males. Non-linear patterns in the relationship between D10 mass and prevalence were also tested but did not significantly improve the model (P > 0.6 for the quadratic and cubic terms).

**Figure 1:**
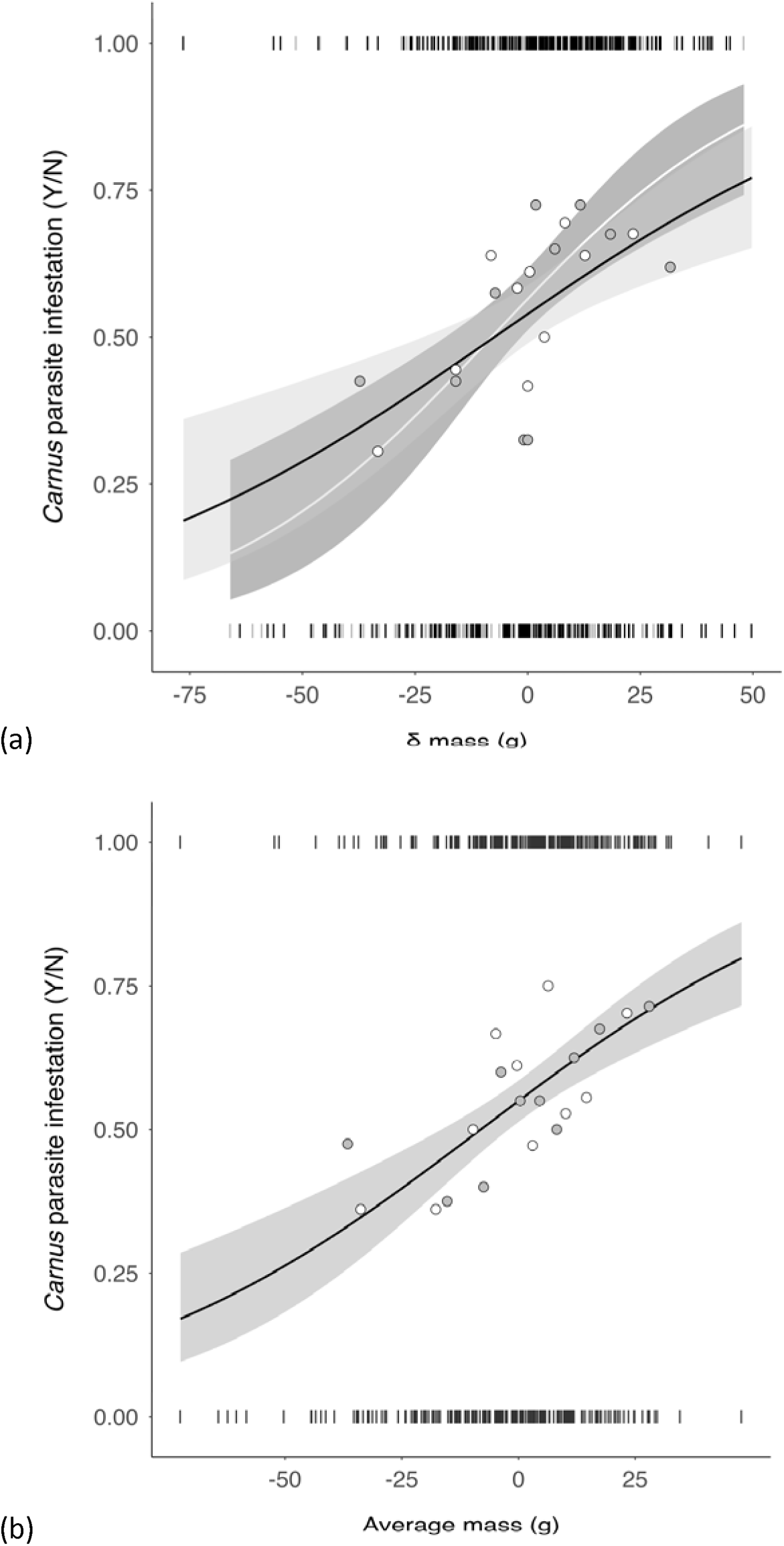
*Carnus* parasite infestation (Y = presence, N = absence) in relation to mass in 10-day-old jackdaw nestlings and regression lines extracted from the model shown in Table 1, with ribbons (shaded areas) representing the 95% confidence interval. (a) Effect of within-brood variation in mass separated by sex: for females, white line and dark ribbon; for males, solid line and light grey ribbon. (b) Effect of between-brood variation in mass. Only one regression line was plotted in panel (b) because the average mass effect did not differ between the sexes (see Table 1). Raw values on *Carnus* prevalence are shown as vertical bars on top and bottom, in black for males and grey for females. Averaged prevalence for mass bins is shown as dots (grey = males, n=402; white = females, n=361).

**Table 1.**
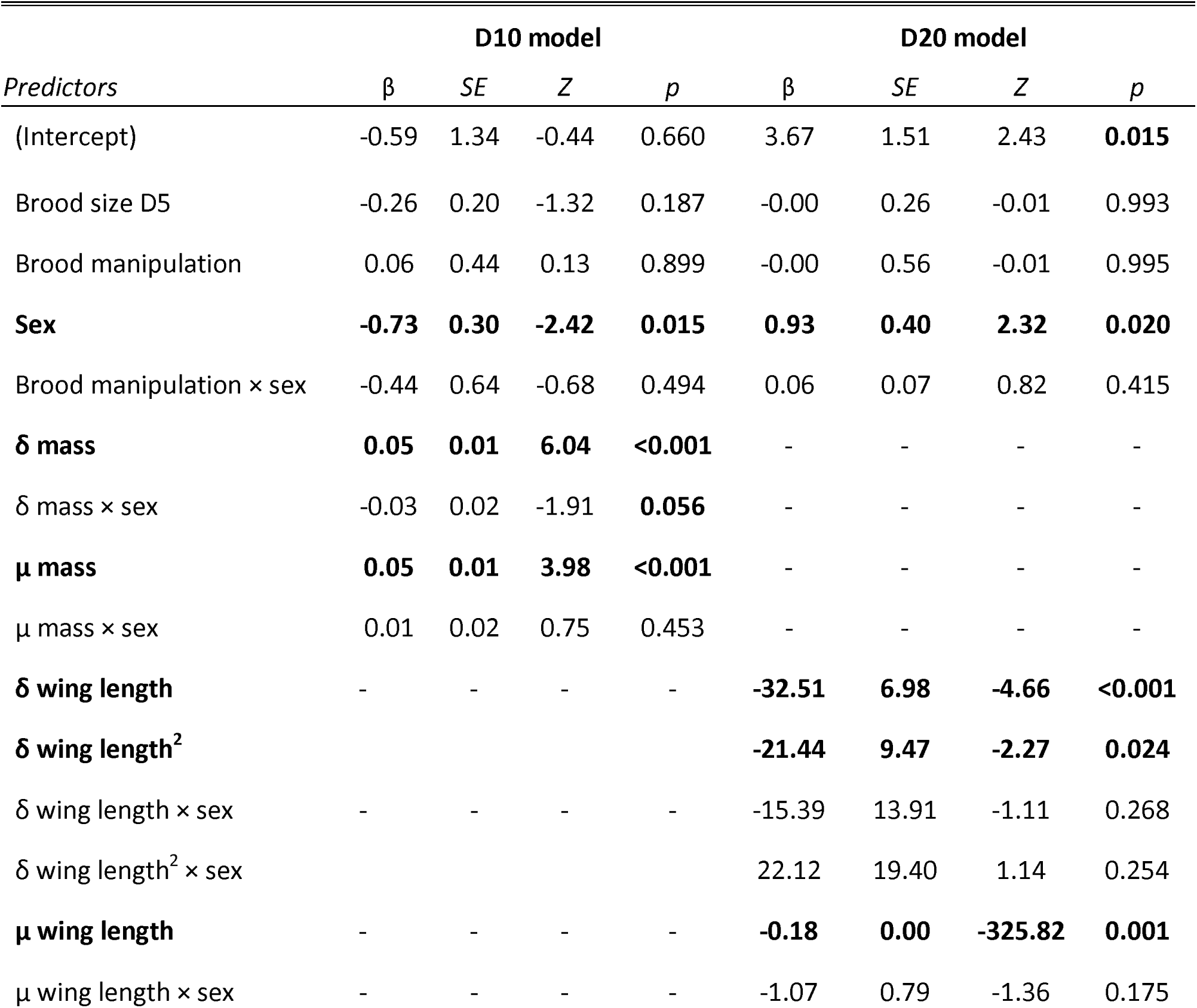
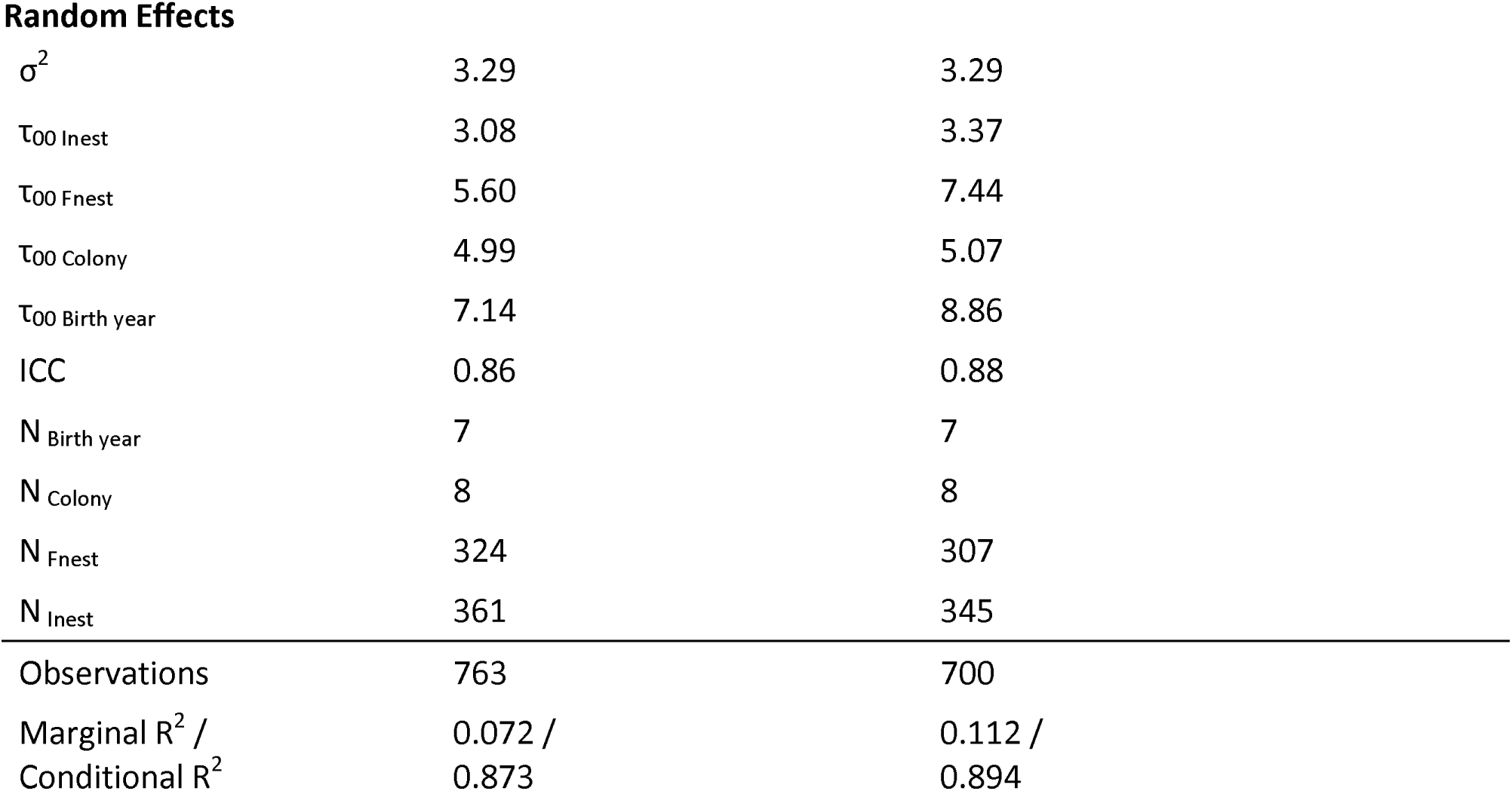
Parameter estimates from a generalised linear mixed model with binomial error distribution on the effect of sex (females coded as base level in all tables), size and brood size manipulation on the likelihood of infection by *Carnus hemapterus*. Note that for mass, within and between-brood effects are separated and that all fixed effects are centred. Average mass and average wing length were centred by brood manipulation category and hence the estimate reflects within category effects. Variable names, parameter estimates (β represent the log odds ratios as extracted from the logistic regression model), statistic (Z) and P – value (p) are in bold when significant. D10 model refers to 10-day-old nestling jackdaws; D20 model: 20-day-old nestlings. Key: SE = standard error, Z = Z-statistic, p = Wald p-value, σ^2^ = residual variance, τ = variance, ICC = interclass correlation coefficient, δ = within-brood effects, μ = between-brood effects, Fnest = foster nest, Inest = incubation nest.

| *Predictors* | **D10 model** |  |  |  | **D20 model** |  |  |  |
| --- | --- | --- | --- | --- | --- | --- | --- | --- |
| | $\beta$ | SE | Z | p | $\beta$ | SE | Z | p |
| (Intercept) | -0.59 | 1.34 | -0.44 | 0.660 | 3.67 | 1.51 | 2.43 | **0.015** |
| Brood size D5 | -0.26 | 0.20 | -1.32 | 0.187 | -0.00 | 0.26 | -0.01 | 0.993 |
| Brood manipulation | 0.06 | 0.44 | 0.13 | 0.899 | -0.00 | 0.56 | -0.01 | 0.995 |
| **Sex** | **-0.73** | **0.30** | **-2.42** | **0.015** | **0.93** | **0.40** | **2.32** | **0.020** |
| Brood manipulation × sex | -0.44 | 0.64 | -0.68 | 0.494 | 0.06 | 0.07 | 0.82 | 0.415 |
| **$\delta$ mass** | **0.05** | **0.01** | **6.04** | **<0.001** | - | - | - | - |
| $\delta$ mass × sex | -0.03 | 0.02 | -1.91 | **0.056** | - | - | - | - |
| **$\mu$ mass** | **0.05** | **0.01** | **3.98** | **<0.001** | - | - | - | - |
| $\mu$ mass × sex | 0.01 | 0.02 | 0.75 | 0.453 | - | - | - | - |
| **$\delta$ wing length** | - | - | - | - | **-32.51** | **6.98** | **-4.66** | **<0.001** |
| **$\delta$ wing length<sup>2</sup>** | - | - | - | - | **-21.44** | **9.47** | **-2.27** | **0.024** |
| $\delta$ wing length × sex | - | - | - | - | -15.39 | 13.91 | -1.11 | 0.268 |
| $\delta$ wing length<sup>2</sup> × sex | - | - | - | - | 22.12 | 19.40 | 1.14 | 0.254 |
| **$\mu$ wing length** | - | - | - | - | **-0.18** | **0.00** | **-325.82** | **0.001** |
| $\mu$ wing length × sex | - | - | - | - | -1.07 | 0.79 | -1.36 | 0.175 |

**Random Effects**
|  |  |  |
| --- | --- | --- |
| $\sigma^2$ | 3.29 | 3.29 |
| $\tau_{00 \text{ Inest}}$ | 3.08 | 3.37 |
| $\tau_{00 \text{ Fnest}}$ | 5.60 | 7.44 |
| $\tau_{00 \text{ Colony}}$ | 4.99 | 5.07 |
| $\tau_{00 \text{ Birth year}}$ | 7.14 | 8.86 |
| ICC | 0.86 | 0.88 |
| $N_{\text{Birth year}}$ | 7 | 7 |
| $N_{\text{Colony}}$ | 8 | 8 |
| $N_{\text{Fnest}}$ | 324 | 307 |
| $N_{\text{Inest}}$ | 361 | 345 |
| Observations | 763 | 700 |
| Marginal $R^2$ / | 0.072 / | 0.112 / |
| Conditional $R^2$ | 0.873 | 0.894 |

On day 20, carnid prevalence was higher than at day 10, with flies recorded on 74% of all nestlings (n = 700). *Carnus* presence on 20-day-old nestlings decreased with increasing wing length within the brood, and this effect was steep as indicated by a significant quadratic effect (Table 1; Fig. 2); note that this effect is opposite to the pattern on day 10, when the heaviest nestlings were more likely to be parasitized. The marginal effect prediction of *Carnus* prevalence was 100% [100%, 100%] when wing length was 1 SD (=7.96 mm) shorter than the mean, 98% [98%, 98%] at the mean wing length, and decreased to 91% [89%, 93%] and 58% [39%, 75%] on nestlings with respectively 1 and 2 SD wing lengths longer than the mean. Between brood variation in wing length was also negatively associated with the likelihood of infection (Table 2), mirroring our findings for the day 10 mass effect (Table 1). As explained in the methods, we used wing length rather than mass in the D20 model, because wing length and mass were strongly correlated (r = 0.80, df = 718, p << 0.001; Fig. S1). Indeed, replacing wing length with mass in the D20 model yields very similar effects although with a slightly lower fit to the data (marginal R^2^ = 0.071, Table S3 vs. marginal R^2^ = 0.112, Table 1).

**Figure 2:**
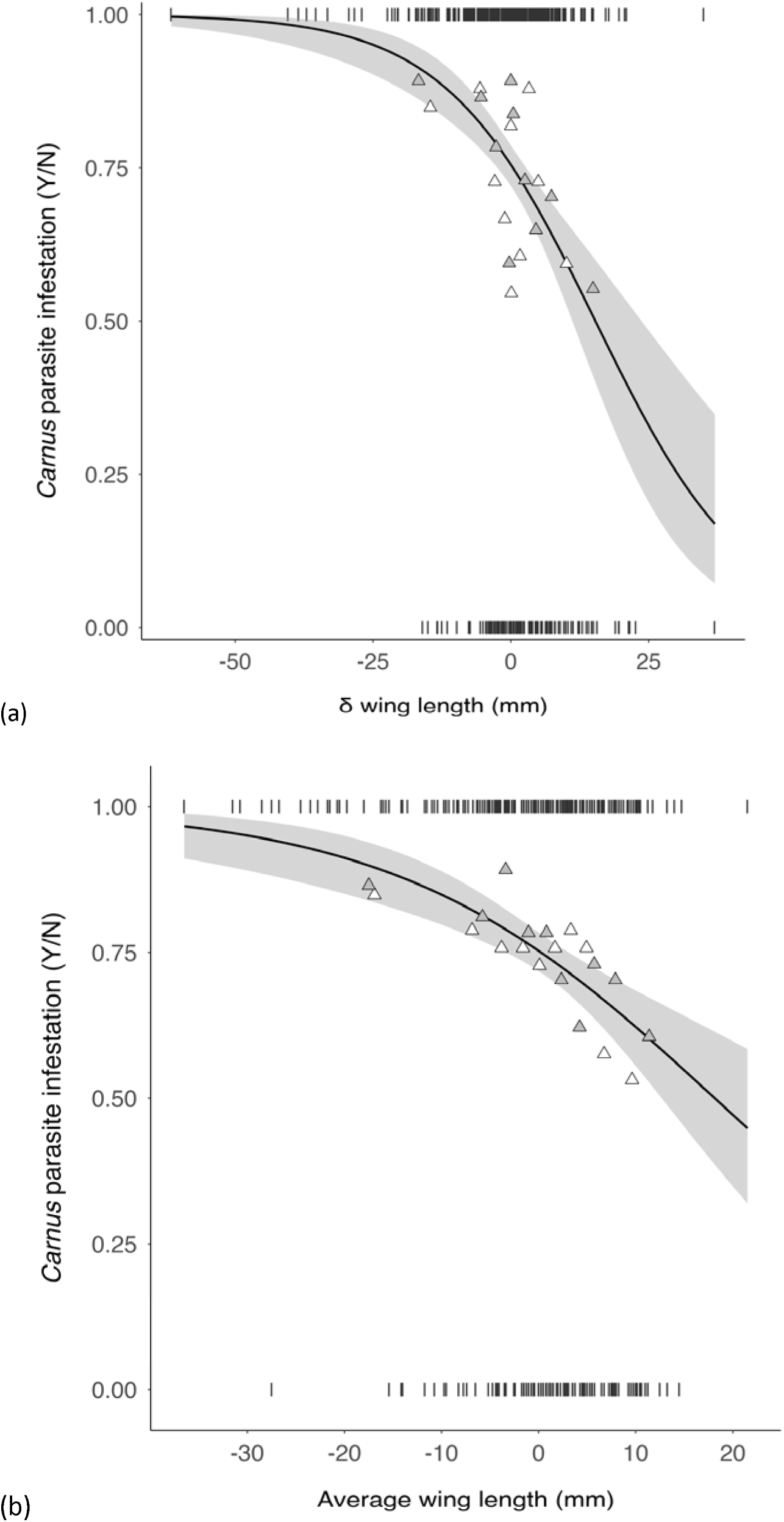
*Carnus* parasite infestation (Y = presence, N = absence) in relation to wing length in 20-day-old jackdaw nestlings and regression lines extracted from the model shown in Table 1, with ribbon (shaded area) representing 95% confidence interval. (a) Effect of within-brood variation in wing length. (b) Effect of between-brood variation in wing length. Only one regression line was plotted because the average mass effect did not differ between the sexes (see Table 1). Raw values on *Carnus* prevalence are shown on top and bottom as vertical bars, in black for males and grey for females. Averaged prevalence for mass bins is shown as triangles (grey = males, n=371; white = females, n=329).

**Table 2.**
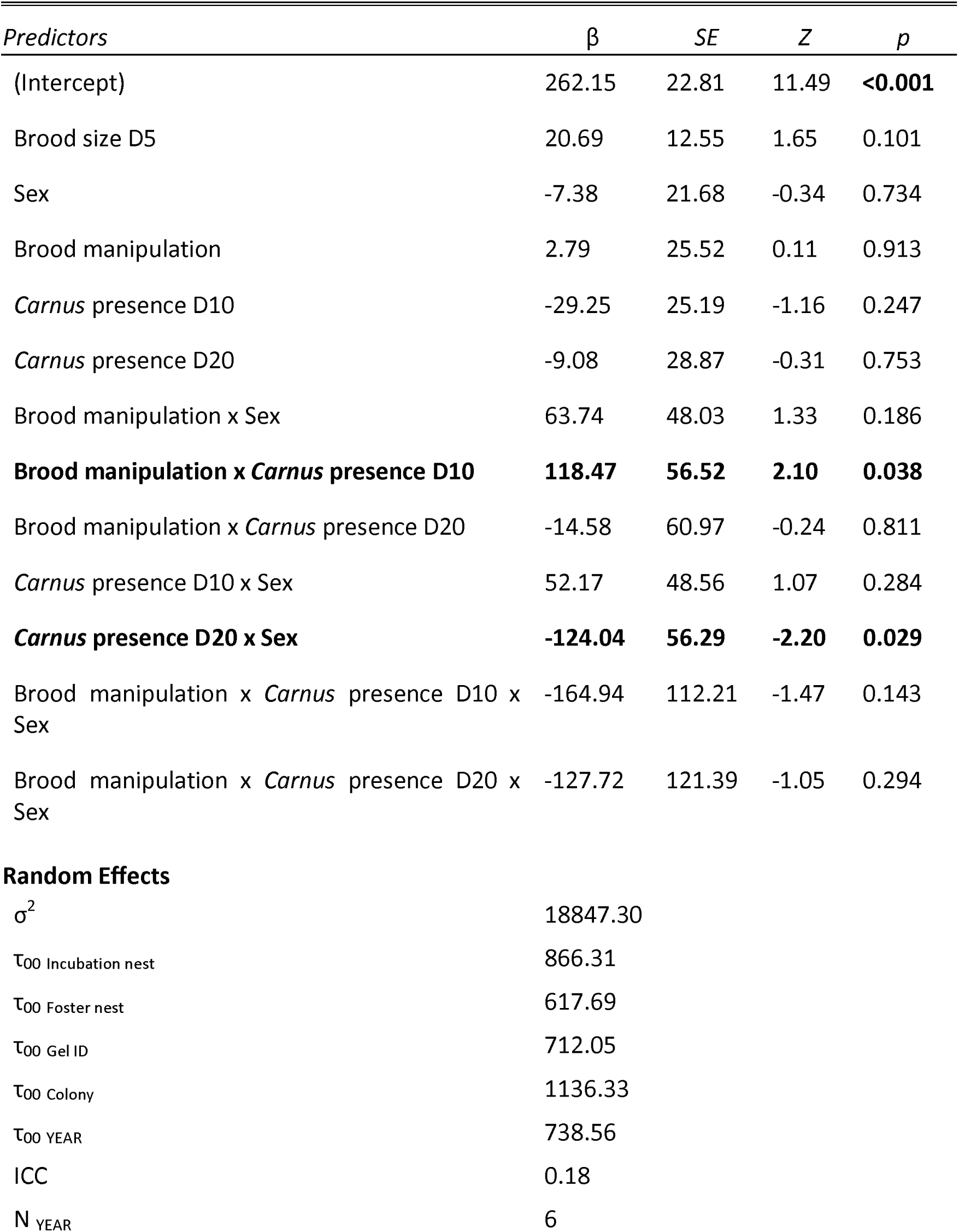
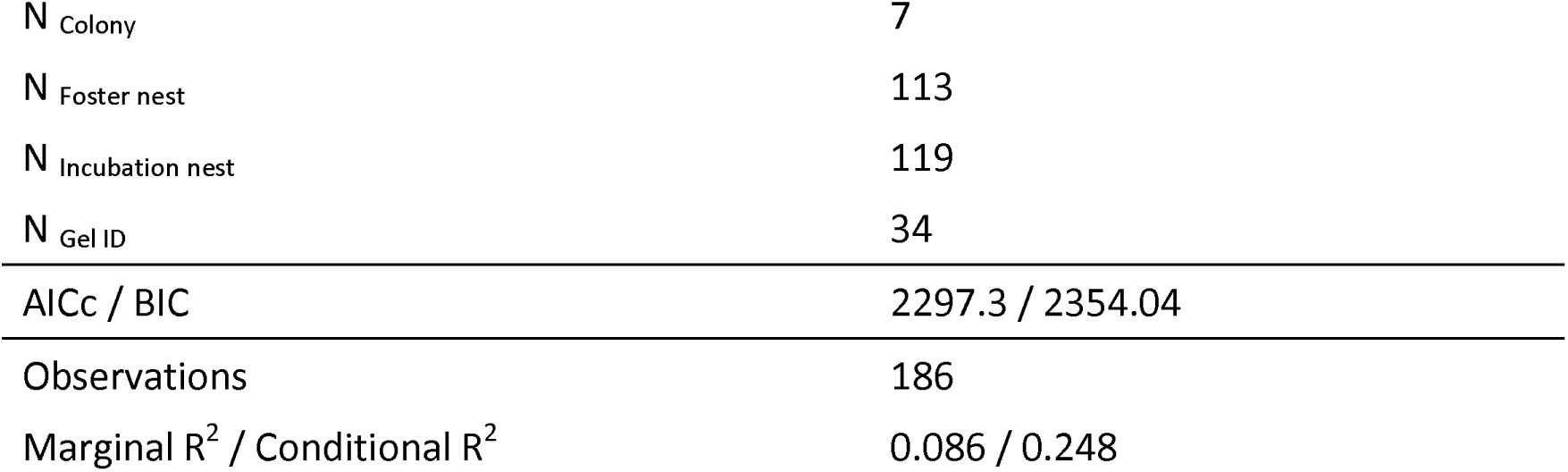
Parameter estimates from a linear mixed model on the effect of *Carnus* presence, sex, and brood size manipulation on telomere shortening in jackdaw nestlings. β = parameter estimate, SE = standard error, Z = Z-statistic, p = Wald p-value, D10 = day 10 of nestling age, D20 = day 20 of nestling age, AICc = corrected Akaike Information Criterion, BIC = Bayesian Information Criterion. Significant fixed effects are in bold.

| *Predictors* | $\beta$ | *SE* | *Z* | *p* |
| --- | --- | --- | --- | --- |
| (Intercept) | 262.15 | 22.81 | 11.49 | **<0.001** |
| Brood size D5 | 20.69 | 12.55 | 1.65 | 0.101 |
| Sex | -7.38 | 21.68 | -0.34 | 0.734 |
| Brood manipulation | 2.79 | 25.52 | 0.11 | 0.913 |
| *Carnus* presence D10 | -29.25 | 25.19 | -1.16 | 0.247 |
| *Carnus* presence D20 | -9.08 | 28.87 | -0.31 | 0.753 |
| Brood manipulation x Sex | 63.74 | 48.03 | 1.33 | 0.186 |
| **Brood manipulation x *Carnus* presence D10** | **118.47** | **56.52** | **2.10** | **0.038** |
| Brood manipulation x *Carnus* presence D20 | -14.58 | 60.97 | -0.24 | 0.811 |
| *Carnus* presence D10 x Sex | 52.17 | 48.56 | 1.07 | 0.284 |
| ***Carnus* presence D20 x Sex** | **-124.04** | **56.29** | **-2.20** | **0.029** |
| Brood manipulation x *Carnus* presence D10 x Sex | -164.94 | 112.21 | -1.47 | 0.143 |
| Brood manipulation x *Carnus* presence D20 x Sex | -127.72 | 121.39 | -1.05 | 0.294 |
| **Random Effects** |  |  |  |  |
| $\sigma^2$ | 18847.30 | | | |
| $\tau_{00}$ Incubation nest | 866.31 | | | |
| $\tau_{00}$ Foster nest | 617.69 | | | |
| $\tau_{00}$ Gel ID | 712.05 | | | |
| $\tau_{00}$ Colony | 1136.33 | | | |
| $\tau_{00}$ YEAR | 738.56 | | | |
| ICC | 0.18 |  |  |  |
| N YEAR | 6 |  |  |  |

|  |  |
| --- | --- |
| N<sub>Colony</sub> | 7 |
| N<sub>Foster nest</sub> | 113 |
| N<sub>Incubation nest</sub> | 119 |
| N<sub>Gel ID</sub> | 34 |
| AICc / BIC | 2297.3 / 2354.04 |
| Observations | 186 |
| Marginal R<sup>2</sup> / Conditional R<sup>2</sup> | 0.086 / 0.248 |

*Carnus* presence on day 20 was negatively related to *Carnus* presence on day 10 (parametric bootstrapped estimate with 1,000 iterations from logistic regression model: β = -3.39, SE = 0.77, Z – value = -4.37, P – value <<0.001). Thus, the contrast between the findings at nestling days 10 and 20, with opposite associations between size and *Carnus* presence, is in line with the negative correlation between *Carnus* presence in early- and late-development: when a focal nestling was not parasitized on day 10, it was more likely that it was parasitized on day 20 and *vice versa*. This shift in carnid presence is further consistent with the strong positive correlation between mass on day 10 and wing length on day 20 (Fig. S1): nestlings that were small on day 10, and hence unlikely to be parasitized, were more likely to be parasitized on day 20 because at that age they were also relatively small.

### Parasite presence and growth effects on telomere shortening

Within individuals, telomere length on days 5 and 30 was strongly correlated (Fig. S2; r = 0.97, n = 184, P < 0.0001). Telomere shortening over the 25-day measurement period was on average 268 ± 23 base pairs (hereafter, bp) ± standard error (Table 2, intercept), or 10.7 bp / day. Testing whether *Carnus* presence early and later during development impacted telomere shortening showed an interaction between brood size manipulation and *Carnus* presence on day 10, and an interaction between sex and *Carnus* presence on day 20 (Table 2). However, these interaction effects were in opposite directions, similar in magnitude (β_Carnus D10 x brood manipulation_ = 118.47 vs. β_Carnus D20 x sex_ = -124.04) and difficult to interpret, given the negative correlation between *Carnus* presence on day 10 vs. day 20. To further investigate these effects, we therefore estimated biologically significant contrasts by manipulation category and sex group. We specifically tested whether being infected only early (vs. late) during development (contrast *Carnus* D10 = 1 - *Carnus* D20 = 0) *vs*. being infected only late during development (contrast *Carnus* D10 = 0 - *Carnus* D20 = 1) affected telomere shortening significantly differently across groups. In females reared in reduced broods, *Carnus* exposure at day 20 resulted in significantly greater telomere shortening compared to exposure at day 10 (Table S4, Δ = −226 ± 77 bp, df = 14, t-ratio = -2.933, P = 0.004, Fig. 3). This effect was weaker and non-significant in enlarged broods. In contrast, males reared in enlarged broods showed a tendency for more telomere loss when they were infected early in development, compared to being infected at a later stage (Table S4, Δ = -93 ± 55 bp, df = 158, t-ratio = 1.703, P = 0.09, Fig. 3). Therefore, despite the main 3-way interaction between *Carnus* presence x brood manipulation x sex being non-significant, our results show biologically significant subgroup patterns indicating that an early *vs.* late parasitic infection during development affects telomere shortening differently in males and females.

**Figure 3:**
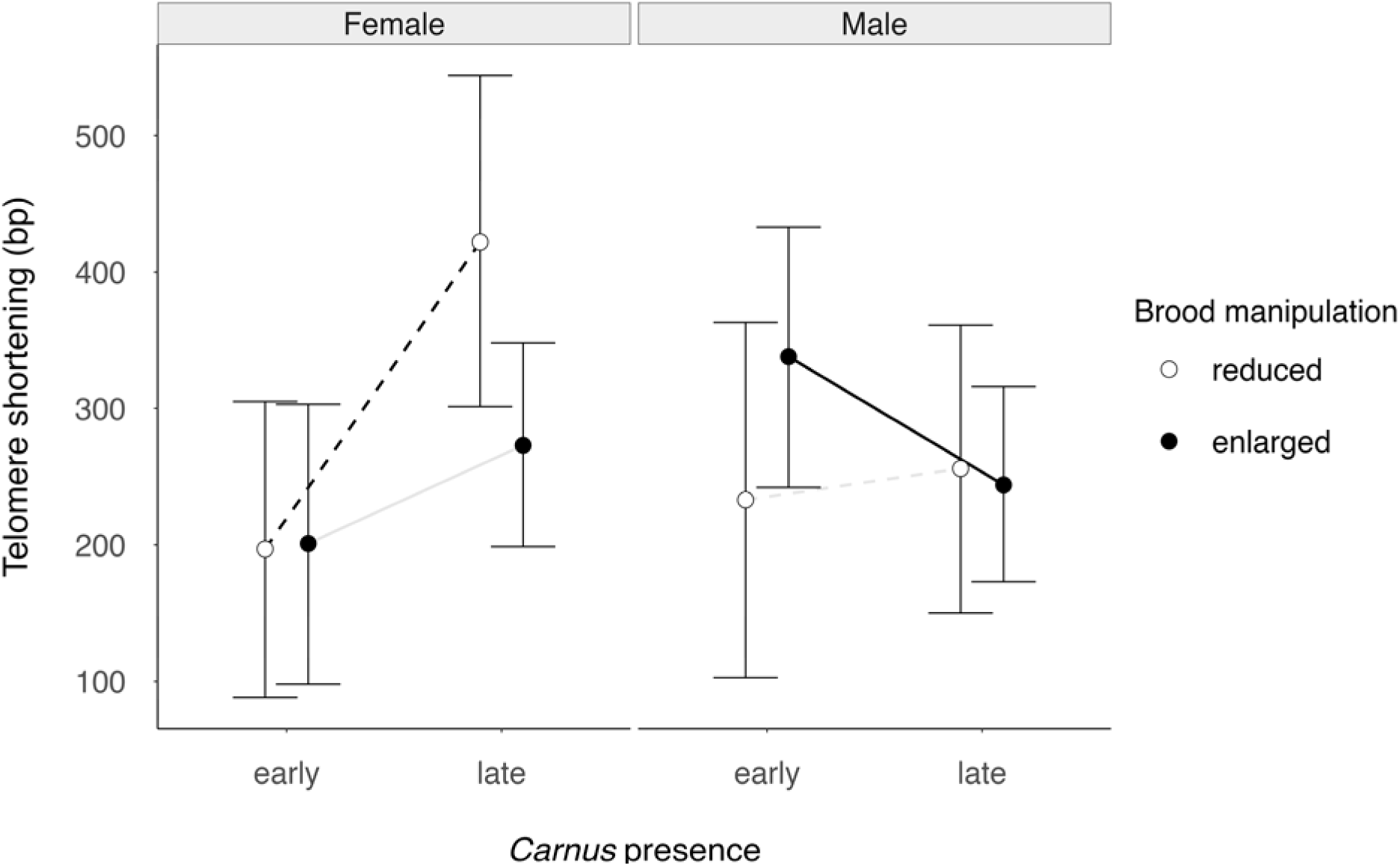
Telomere shortening in relation to *Carnus* presence early vs. late in development in jackdaw nestlings. Data points (estimated marginal means) and confidence interval (bars) are shown separately for females (left panel) and males (right panel) and for brood manipulation categories. Note that the dashed (reduced broods) and solid lines (enlarged broods) join the means from the early and late-infected nestlings only for representation purposes. Lines appear in black when the contrast was significant (females, reduced group, dashed line) or marginally non-significant (males, enlarged group, solid line), in grey when not significant. Key: bp = base pairs, early = *Carnus* parasites were present when nestlings were 10 days old and not present on day 20, late = *Carnus* present only when nestlings were 20 days old.

When evaluating whether mass on day 10 or wing length on day 20 (proxies of *Carnus* prevalence), explained telomere shortening over the developmental period, we found that telomere shortening was not associated with mass on day 10 or wing length on day 20 (Table 3), despite the size associations with *Carnus* infection. There was, however, a sex-specific association between growth and telomere shortening, as indicated by a significant interaction between day 10 mass and sex. Males lost on average 7±3 more base pairs per gram increase of day 10 mass than females, in which day 10 mass was not significantly associated with telomere shortening (Table 3, Fig. 4). This is in accordance with the finding that 20-day old male nestlings that carried parasites (and were not infected on day 10) lost marginally non-significantly fewer telomere base pairs than females (Table 2).

**Figure 4:**
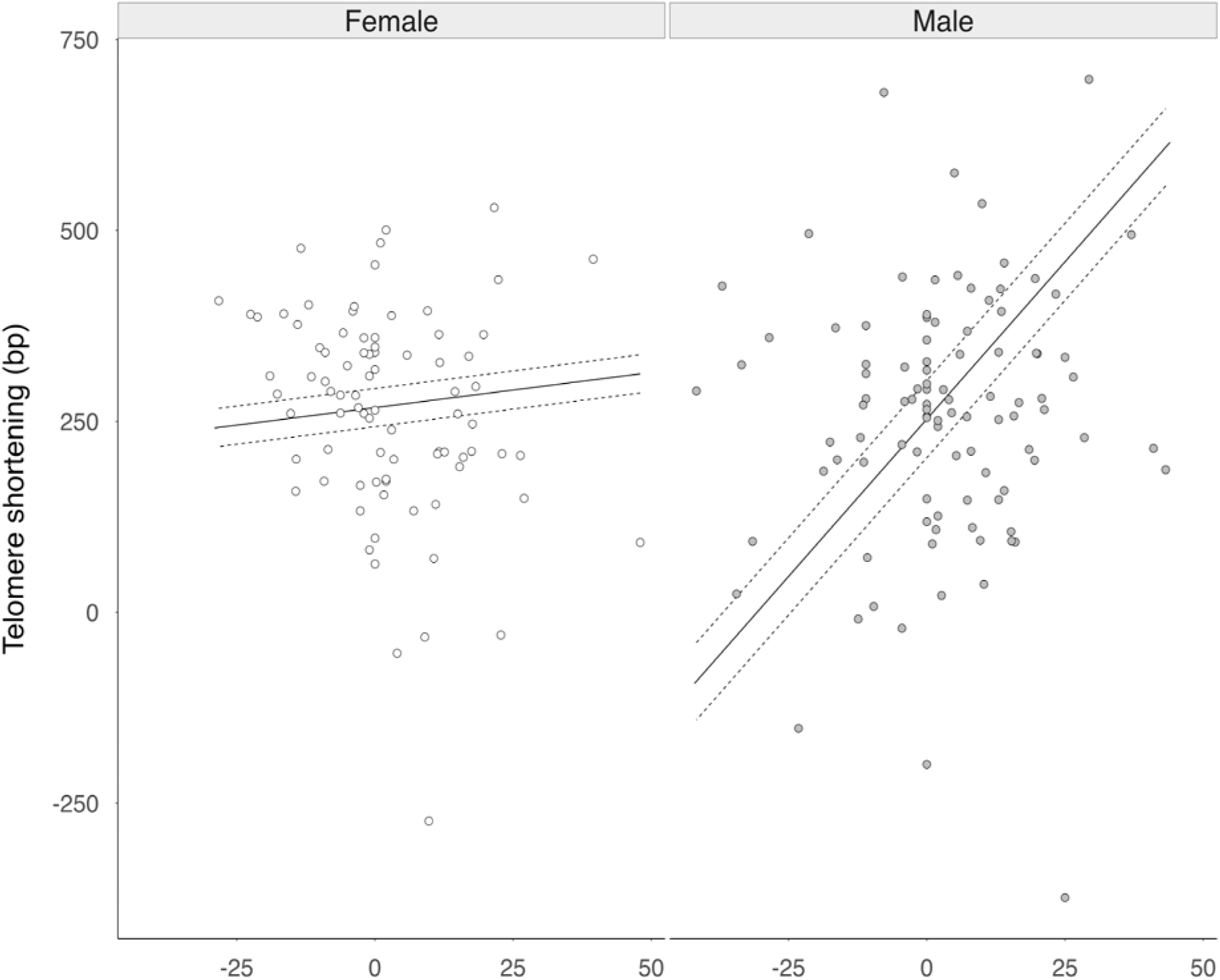
Telomere shortening in relation to mass and sex in 10-day-old jackdaw nestlings. Regression lines were extracted from the model in Table 2, with ribbons (dashed lines) representing standard error. Raw values are shown as dots (grey = males, n=100; white = females, n=84). Key: bp = base pairs, δ mass D10 = within-brood effects of mass on 10-day-old nestlings, f = females, m = males.

**Table 3.**
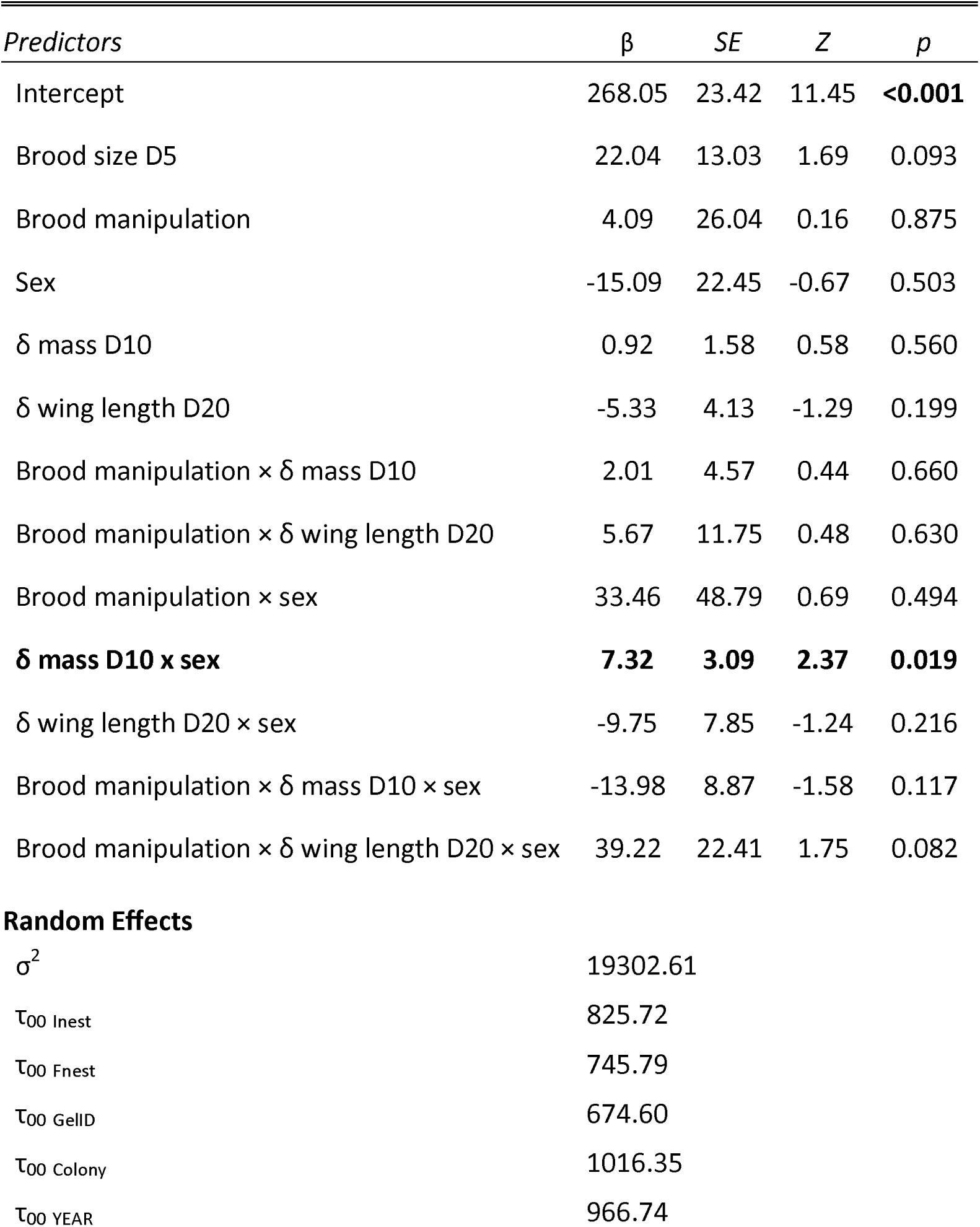
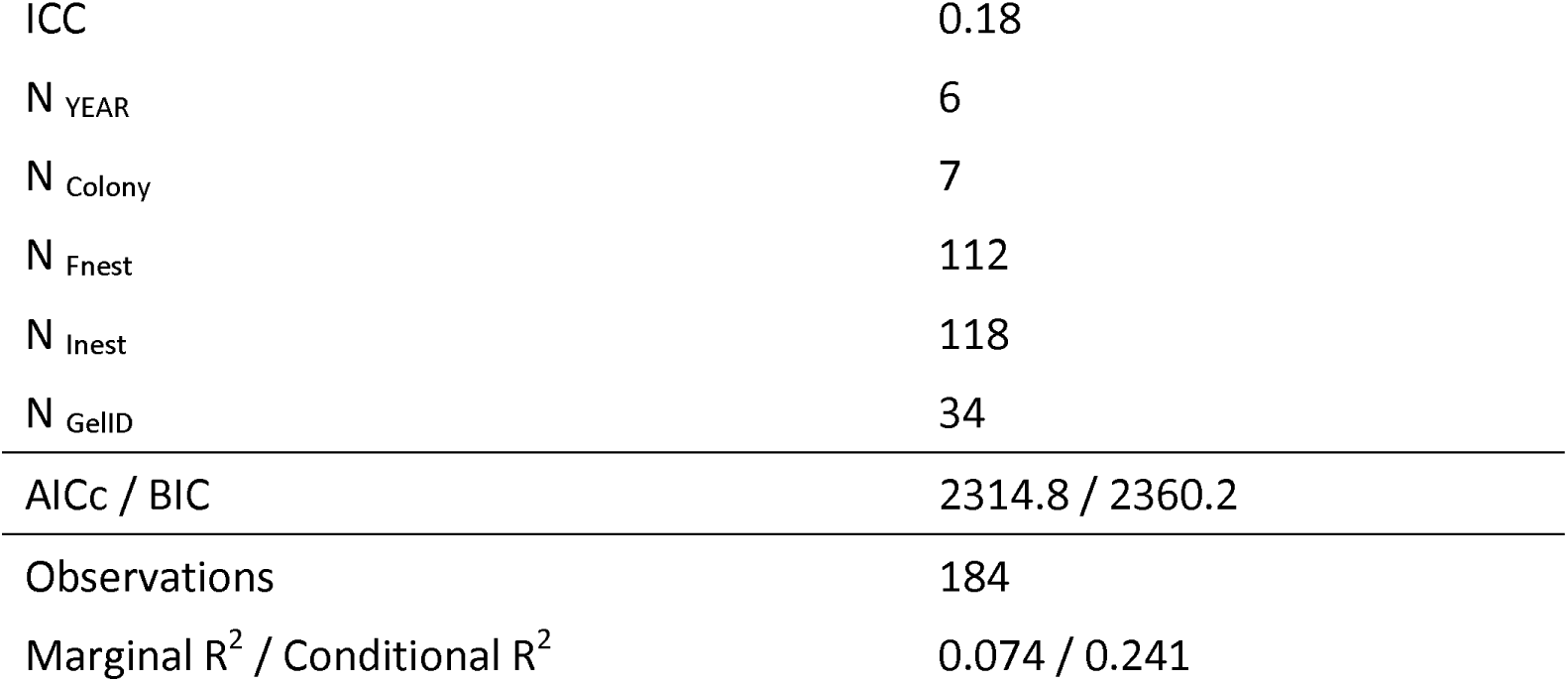
Parameter estimates from a linear mixed model on the effects of growth, sex and brood size manipulation on telomere shortening in jackdaw nestlings. Note that all fixed effects are centred. Variable names, parameter estimates (β), statistics (Z) and Wald p – values (p) are in bold when significant. D10 refers to 10-day-old nestling jackdaws; and D20 to 20-day-old nestlings. Key: SE = standard error, σ^2^ = residual variance, τ = variance, ICC = interclass correlation coefficient, δ = within-brood effects, Fnest = foster nest, Inest = incubation nest, AICc = corrected Akaike Information Criterion, BIC = Bayesian Information Criterion.

## DISCUSSION

We investigated parasite presence and telomere shortening during development in a wild population of jackdaws. Our main findings were that the carnid fly *Carnus hemapterus* infection pattern varied with respect to nestling sex and size in an age dependent way, and that the timing of the infection during the nestling period may affect telomere shortening differently in males and females. A major inference from our results is that nestling infection patterns and associated effects can be age dependent. For example, males were less likely to be infected than females on day 10, but more likely to be infected on day 20 than females, independent of their size. In this setting, estimating parasite prevalence at a single timepoint during development is likely to yield a biased picture of the infection patterns and consequences.

The ‘tasty-chick’ hypothesis (Christe et al. 1998) states that the less immunocompetent later-hatched nestlings act as ‘bait’ for parasites, and consequently carry higher parasite loads. However, (Garrido-Bautista 2025), reviewing tests of the tasty-chick hypothesis, concluded that “Most studies show that ectoparasites tend to aggregate on larger and high-quality nestlings within broods, rather than in the late-hatched nestlings.” In line with this conclusion, we observed higher prevalence of *Carnus* infection on the larger nestlings early in the nestling period (day 10), which also in jackdaws are the older nestlings in the brood. Earlier studies have focused primarily on variation in parasite prevalence within broods (Garrido-Bautista 2025), but here we simultaneously investigated how *Carnus* prevalence associated with variation in mass / size within *and* between broods. Interestingly, we found that associations between mass and *Carnus* prevalence within and between broods were indistinguishable. This indicates that, for example, the positive association between day 10 mass and *Carnus* prevalence within broods is not only due to parasites in a nest selecting larger nestlings to feed from within a brood, but also that a higher proportion of nestlings was infected when the average day 10 mass in a brood was higher. Whether this was due to higher *Carnus* fly numbers present in nest boxes where day 10 mass was high, or due to more carnid flies leaving the nest material to forage on the nestlings when day 10 mass was high, we cannot evaluate with the present data set.

However, the opposite pattern emerged late in the nestling period (day 20); as also observed in barn owl nestlings (Roulin et al. 2003, 2008). Such a change in infection pattern throughout development can be interpreted as the outcome of ectoparasite preference, with *Carnus* parasites preferentially feeding on nestlings in a specific size range. In this scenario, the larger nestlings on day 10 are closer to the desired size, while on day 20 the smaller nestlings are closer to the desired size. Alternatively, but not exclusively, on day 20, the flies may have infected the smallest and lightest nestlings because these are less densely covered by feathers than their more developed / older siblings (Liker et al., 2011; A. Roulin et al., 2008).

Infection patterns are the outcome of host-parasite interactions, shaped by the host’s quality vs. its vulnerability to infection (Attwood 2025). This raises the question to what extent the infection patterns we observed reflect host behaviours rather than parasite preference. Anti-parasitic behaviours such as grooming have been reported in nestlings (Clayton 1991) and, in some species, parents rearing infested nestlings engage more frequently in ectoparasite removal (e.g. in several passerine species (Cantarero et al. 2013b)). We do not know from what age jackdaw nestlings can groom themselves, but given the agility and motor skills this requires we consider it unlikely before day 20. However, anecdotal observations have confirmed that females groom their nestlings (Verhulst, unpublished); whether they do this selectively with respect to mass or size is, at present, unknown. Medication experiments that include within-brood parasite removal in combination with detailed behavioural observations should shed light on these hypotheses.

Regarding potential detrimental effects of the parasitic infection on jackdaws, we previously showed that carnid fly infections of a brood at any point during development (exacerbated by sibling competition) accelerated telomere shortening but did not affect body mass gain (Badás et al., 2023). Here, examining infection patterns among nestlings within infected broods only, we found the timing of the

*Carnus* infection to have sex-specific effects on telomere shortening. Female nestlings suffered more telomere shortening when infected with *Carnus* later in development, and this effect was apparent in the reduced broods only. In contrast, males that grew heavier than their siblings up to day 10, and thus were more likely to be infected earlier in development, lost more telomere base pairs, and this effect increased with mass. Faster growth could pose an additional challenge to telomere shortening under infection. Indeed, it has been associated with increased telomere loss across taxa, e.g., in several bird species (Vedder et al. 2018; Salmón et al. 2021; Kärkkäinen et al. 2022), in fish (Pauliny et al. 2015) and in lizards (Zhang et al. 2023), but not in amphibians (Burraco et al. 2017)).

Sex dependent parasite effects on telomere attrition have previously been observed in mice *Mus musculus* (Ilmonen et al. 2008) and birds (Tschirren et al. 2021). Two mutually non-exclusive explanations could be behind the sex-specific effects on telomere shortening: first, there might be sex differences in parasite susceptibility and/or immunity (Tschirren et al. 2003; Restif and Amos 2010; Romano et al. 2011; Driessen et al. 2022). And a second, not mutually exclusive, hypothesis is that sexes differentially allocate resources to counteract telomere shortening (i.e. telomere repair). A different allocation response to infection is predicted when the relationship between early-life telomere length and fitness prospects is sex-dependent (Tschirren et al. 2021). Male jackdaws gain significantly more weight than females over the 25-day developmental period (Badás et al. 2023), which may indicate that males allocate more resources to body mass gain or that males and females show different growth curves that may influence the timing of infection. Indeed, male jackdaws are significantly heavier than females at fledgling ((Arnold and Griffiths 2003); also in the present study: linear mixed model of sex effects on fledgling mass, β estimate for males = 19.59 g, SE = 2.33, df = 141.97, t-value = 8.41, P<0.0001). However, as we currently do not know (i) the exact carnid loads across siblings and sexes, and/or (ii) whether early-life telomere shortening has sex-specific effects on fitness, both hypotheses remain speculative at this stage.

In our previous study (Badás et al. 2023), we highlighted that sibling competition and ectoparasite infection synergistically impacted telomere shortening (Badás et al. 2023). We also suggested that resources may be canalised to body mass gain (Boonekamp et al. 2018), but this hypothesis requires further investigation. Here, we found additional support for the ‘reverse Matthew effect’, by which individuals in a poor state ended up in an even worse state (increased telomere loss) after facing not only environmental (ectoparasite infection) but also internal stressors (fast growth in males). In the present study, we pave the way for disentangling the particular over-burdening developmental conditions that could prevent canalisation and thus impact fitness later in life (e.g., for males, investing on rapid early-growth under the pressure of infection). Challenging conditions have been suggested to impair the ability to activate physiological compensatory mechanisms later in life in individuals at the ‘fast end’ of the pace-of-life continuum, i.e., individuals that grow faster, show higher metabolic rates and/or earlier reproduction ((Polverino et al. 2018); for a review, (Badás et al. 2025)). We thus highlight that parasitic infections are dynamic processes that may exacerbate external challenges (e.g. sibling competition) and interact with internal conditions (e.g., sex, genotype) differently throughout development. Since telomere shortening in early-life has been proposed as a robust biological marker of somatic redundancy (Boonekamp et al. 2013) or survival prospects (Boonekamp et al. 2014), future studies that measure telomere shortening also throughout adulthood will allow us to extract robust conclusions on the fitness consequences of facing such over-burdening developmental stressors.

## ACKNOWLEDGEMENTS

We thank Martijn Salomons, the late Cor Dijkstra and many students that contributed to the fieldwork.

## Ethics approval

Fieldwork was approved by the University of Groningen animal ethics committee under license numbers (4071, 5871, 6832-A & 6832-B) before conducting research.

## Consent to participate

N/A

## Consent for publication

N/A

## Availability of data and material

Data used for this manuscript will be available upon acceptance, in Zenodo, digital repository with DOI.

## Code availability

Code used for this manuscript will be available upon acceptance, in Zenodo, digital repository with DOI and in https://github.com/elisa-P-badas.

## Author contributions

The study was conceived by EPB and SV. Samples were collected by CB, JB, and SV. Telomere analyses were performed by CB, EM. Statistical analyses were conducted by EPB with input from SV. The paper was written by EPB with extensive input from SV and additional comments from all authors. All authors gave final approval for publication.

