## Supplemental tables and figures for "Growing despite ectoparasite infection: sex-specific effects on telomere shortening in a free-living corvid"

**Electronic supplementary material for:**

**Table of Contents:**

| **Supplementary tables** | Pages 2 – 5 |
| --- | --- |
| **Supplementary figures** | Pages 6 – 7 |

**Table S1.** Parameter estimates from the logistic regression on parasite presence at day 10 in nestling jackdaws (including infected and not-infected broods throughout development). Model 1 includes year as random effect, Model 2 as fixed effect, Model 3 shows the difference between the within and between-broods effects, which justifies separating mass in the within- and between-broods terms. Terms in brackets for factors represent the reference level. β = parameter estimate, SE = standard error, Z = Z statistic, P-value = Wald P-value, X̄ mass = average mass. Significant fixed effects are in bold.

| **Parameter (reference level)** | **β** | **SE** | **Z** | **P** |
| --- | --- | --- | --- | --- |
| ***Model 1 (N = 1319 nestlings, 440 nests)*** |  |  |  |  |
| Intercept | -2.305 | 1.313 | -1.756 | 0.0791. |
| **Mass** | **0.855** | **0.149** | **5.739** | **<0.0001****** |
| Brood size on day 5 | 0.123 | 0.145 | 0.851 | 0.395 |
| Sex (female) |  |  |  |  |
| Male | -0.289 | 0.236 | -1.222 | 0.2218 |
| Brood manipulation (reduced) |  |  |  |  |
| Enlarged | -0.037 | 0.241 | -0.155 | 0.8771 |
| ***Model 2 (N = 1319 nestlings, 440 nests)*** |  |  |  |  |
| Intercept | -12.92 | 1.697 | -7.613 | <0.0001**** |
| Mass | 0.666 | 0.390 | 1.710 | <0.0872. |
| Brood size on day 5 | 0.038 | 0.317 | 0.121 | 0.9033 |
| Sex (female) |  |  |  |  |
| Male | -0.139 | 0.600 | -0.231 | 0.8171 |
| Brood manipulation (reduced) |  |  |  |  |
| Enlarged | 0.013 | 0.600 | 0.021 | 0.9830 |
| Year (2012) |  |  |  |  |
| 2013 | -1.796 | 2.355 | -0.763 | 0.4456 |
| 2014 | -0.460 | 1.730 | -0.266 | 0.7903 |
| **2015** | **21.752** | **1.849** | **11.764** | **<0.0001****** |
| 2016 | 0.456 | 1.684 | 0.271 | 0.7864 |
| 2017 | 0.739 | 1.680 | 0.440 | 0.6602 |
| 2018 | 0.007 | 1.715 | 0.004 | 0.9966 |
| ***Model 3 (N = 1310 nestlings, 437 nests)*** |  |  |  |  |
| Intercept | -3.875 | 1.699 | -2.281 | <0.05* |
| **Mass (within-broods effect)** | **0.044** | **0.007** | **6.035** | **<0.0001****** |
| **X̄ mass (within- vs. between-broods difference)** | **-0.031** | **0.011** | **-2.817** | **<0.01**** |
| Brood size on day 5 | 0.090 | 0.146 | 0.617 | 0.537 |
| Sex (female) |  |  |  |  |
| Male | -0.274 | 0.238 | -1.148 | 0.251 |
| Brood manipulation (reduced) |  |  |  |  |
| Enlarged | -0.092 | 0.244 | -0.380 | 0.704 |

Random effect estimates (variance, Model 1): Nestling ID nested in Foster Nest (2.152),

Foster nest (5.967), Colony (4.989), Year (6.209); Random effect estimates (variance, Model 2):

Nestling ID nested in Foster Nest (1600.531), Foster nest (64.813), Colony (4.806); Random

effect estimates (variance, Model 3): Nestling ID nested in Foster Nest (2.222),

Foster nest (5.878), Colony (4.977), Year (6.295)

**Table S2.** Parameter estimates from the logistic regression on parasite presence at day 20 in nestling jackdaws (including infected and not-infected broods throughout development). Model 1 includes year as random effect, Model 2 as fixed effect, Model 3 shows the difference between the within and between-nest effects. Terms in brackets for factors represent the reference level. β = parameter estimate, SE = standard error, Z = Z statistic, P-value = Wald P-value, X̄ mass = average mass. Significant fixed effects are in bold.

| **Parameter (reference level)** | **β** | **SE** | **Z** | **P** |
| --- | --- | --- | --- | --- |
| ***Model 1 (N = 1319 nestlings, 440 nests)*** |  |  |  |  |
| Intercept | -2.305 | 1.313 | -1.756 | 0.0791. |
| **Mass** | **0.855** | **0.149** | **5.739** | **<0.0001****** |
| Brood size on day 5 | 0.123 | 0.145 | 0.851 | 0.395 |
| Sex (female) |  |  |  |  |
| Male | -0.289 | 0.236 | -1.222 | 0.2218 |
| Brood manipulation (reduced) |  |  |  |  |
| Enlarged | -0.037 | 0.241 | -0.155 | 0.8771 |
| ***Model 2 (N = 1319 nestlings, 440 nests)*** |  |  |  |  |
| Intercept | -12.92 | 1.697 | -7.613 | <0.0001**** |
| Mass | 0.666 | 0.390 | 1.710 | <0.0872. |
| Brood size on day 5 | 0.038 | 0.317 | 0.121 | 0.9033 |
| Sex (female) |  |  |  |  |
| Male | -0.139 | 0.600 | -0.231 | 0.8171 |
| Brood manipulation (reduced) |  |  |  |  |
| Enlarged | 0.013 | 0.600 | 0.021 | 0.9830 |
| Year (2012) |  |  |  |  |
| 2013 | -1.796 | 2.355 | -0.763 | 0.4456 |
| 2014 | -0.460 | 1.730 | -0.266 | 0.7903 |
| **2015** | **21.752** | **1.849** | **11.764** | **<0.0001****** |
| 2016 | 0.456 | 1.684 | 0.271 | 0.7864 |
| 2017 | 0.739 | 1.680 | 0.440 | 0.6602 |
| 2018 | 0.007 | 1.715 | 0.004 | 0.9966 |
| ***Model 3 (N = 1310 nestlings, 437 nests)*** |  |  |  |  |
| Intercept | -3.875 | 1.699 | -2.281 | <0.05* |
| **Mass (within-nest effect)** | **0.044** | **0.007** | **6.035** | **<0.0001****** |
| **X̄ mass (within- vs. between-nests difference)** | **-0.031** | **0.011** | **-2.817** | **<0.01**** |
| Brood size on day 5 | 0.090 | 0.146 | 0.617 | 0.537 |
| Sex (female) |  |  |  |  |
| Male | -0.274 | 0.238 | -1.148 | 0.251 |
| Brood manipulation (reduced) |  |  |  |  |
| Enlarged | -0.092 | 0.244 | -0.380 | 0.704 |

Random effect estimates (variance, Model 1): Nestling ID nested in Foster Nest (2.152),

Foster nest (5.967), Colony (4.989), Year (6.209); Random effect estimates (variance, Model 2):

Nestling ID nested in Foster Nest (1600.531), Foster nest (64.813), Colony (4.806); Random

effect estimates (variance, Model 3): Nestling ID nested in Foster Nest (2.222),

Foster nest (5.878), Colony (4.977), Year (6.295)

**Table S3.** Parameter estimates from the generalised linear mixed model with binomial error distribution on the effects of mass, brood manipulation and sex on *Carnus* presence at day 20 in nestling jackdaws. β = parameter estimate, SE = standard error, Z = Z statistic, P-value = Wald P-value, δ mass = within-individual mass effect, µ mass = between individual mass effect. Significant fixed effects are in bold.

| *Predictors* | β | *SE* | *Z* | *p* |
| --- | --- | --- | --- | --- |
| (Intercept) | 3.67 | 1.52 | 2.42 | **0.016** |
| Brood manipulation | -0.03 | 0.51 | -0.06 | 0.952 |
| Brood size D5 | 0.03 | 0.24 | 0.11 | 0.909 |
| **Sex** | **1.41** | **0.42** | **3.35** | **0.001** |
| **δ mass** | **-0.06** | **0.01** | **-4.57** | **<0.001** |
| **μ mass** | **-0.03** | **0.01** | **-1.98** | **0.048** |
| δ mass × sex | -0.05 | 0.03 | -1.95 | 0.051 |
| μ mass x sex | -0.03 | 0.02 | -1.61 | 0.108 |
| Brood manipulation x sex | -0.82 | 0.75 | -1.11 | **0.268** |
| **Random Effects** | | | | |
| σ^2^ | 3.29 | | | |
| τ_00_ _Incubation nest_ | 3.15 | | | |
| τ_00_ _Foster nest_ | 5.98 | | | |
| τ_00_ _Colony_ | 5.14 | | | |
| τ_00_ _YEAR_ | 9.16 | | | |
| ICC | 0.88 | | | |
| N _YEAR_ | 7 | | | |
| N _Colony_ | 8 | | | |
| N _Foster nest_ | 309 | | | |
| N _Incubation nest_ | 346 | | | |
| Observations | 705 | | | |
| Marginal R^2^ / Conditional R^2^ | 0.071 / 0.886 | | | |

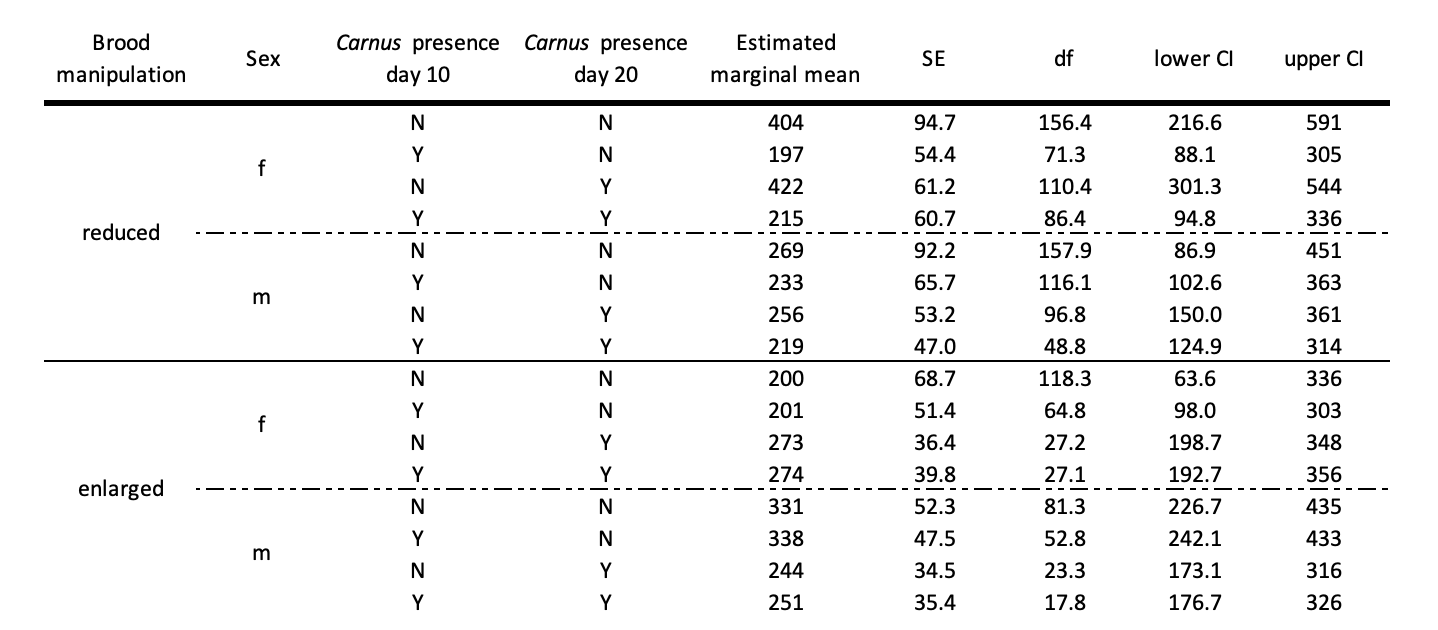
**Table S4:** Estimated marginal means for telomere shortening over the 25-day developmental period in relation to *Carnus* presence, sex and brood manipulation. Key: N = no parasites detected, Y = parasites detected, f = female, m = male, SE = standard error, CI = confidence interval.

**Figure S1:** The association between the within-brood mass when jackdaw nestlings were 10 days old and wing length at 20 days old. Raw data points are shown. Raw data, jittered for clarity on the amount of data, are shown. The fitted line represents the linear regression and 95% confidence intervals are shown as shaded areas.

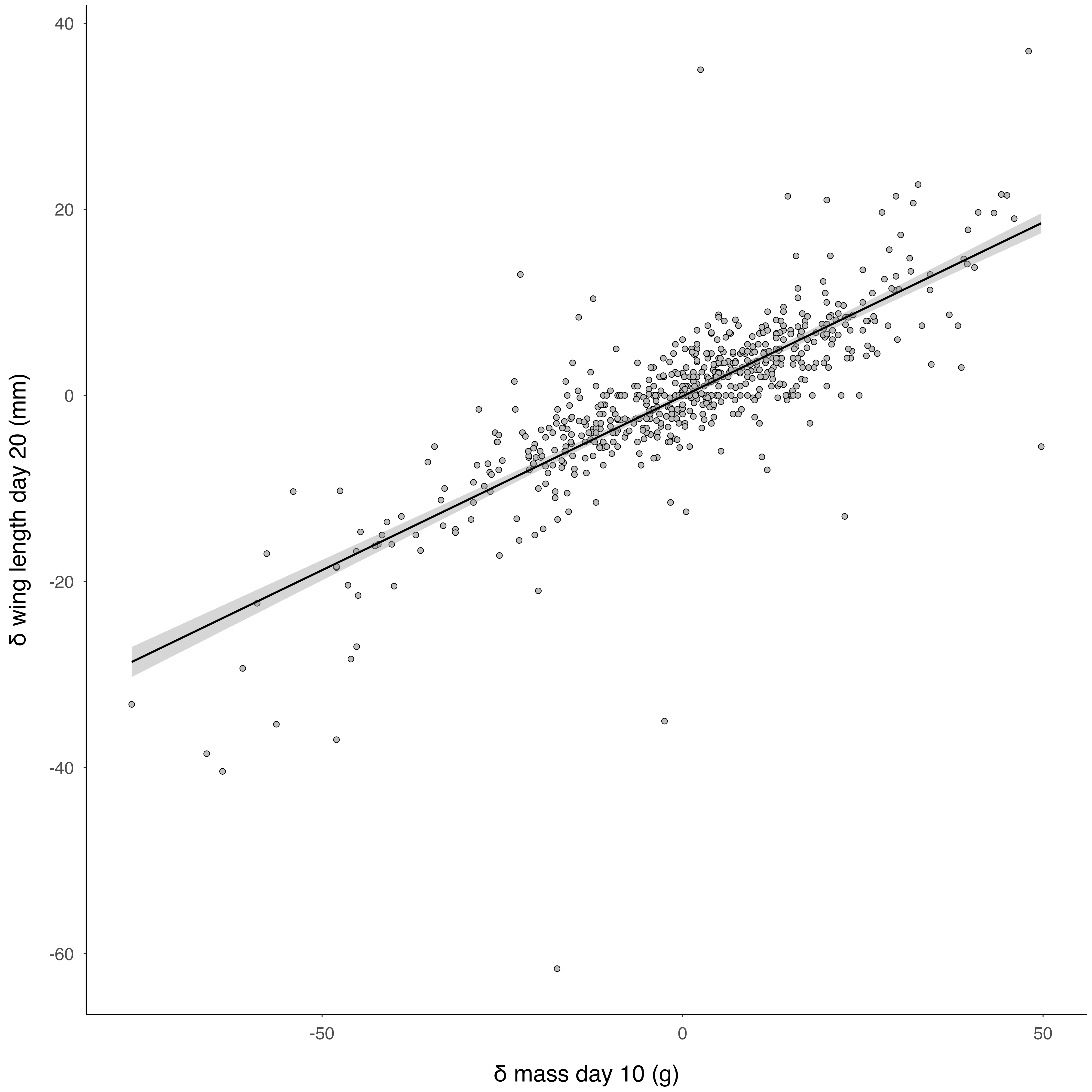

**Figure S2:** Within-individual differences in telomere length in wild jackdaws as shown by the correlation between telomere length at day 5 and day 30. The red diagonal line represents no difference (no change in telomere length) and further points below the line indicate more telomere shortening.

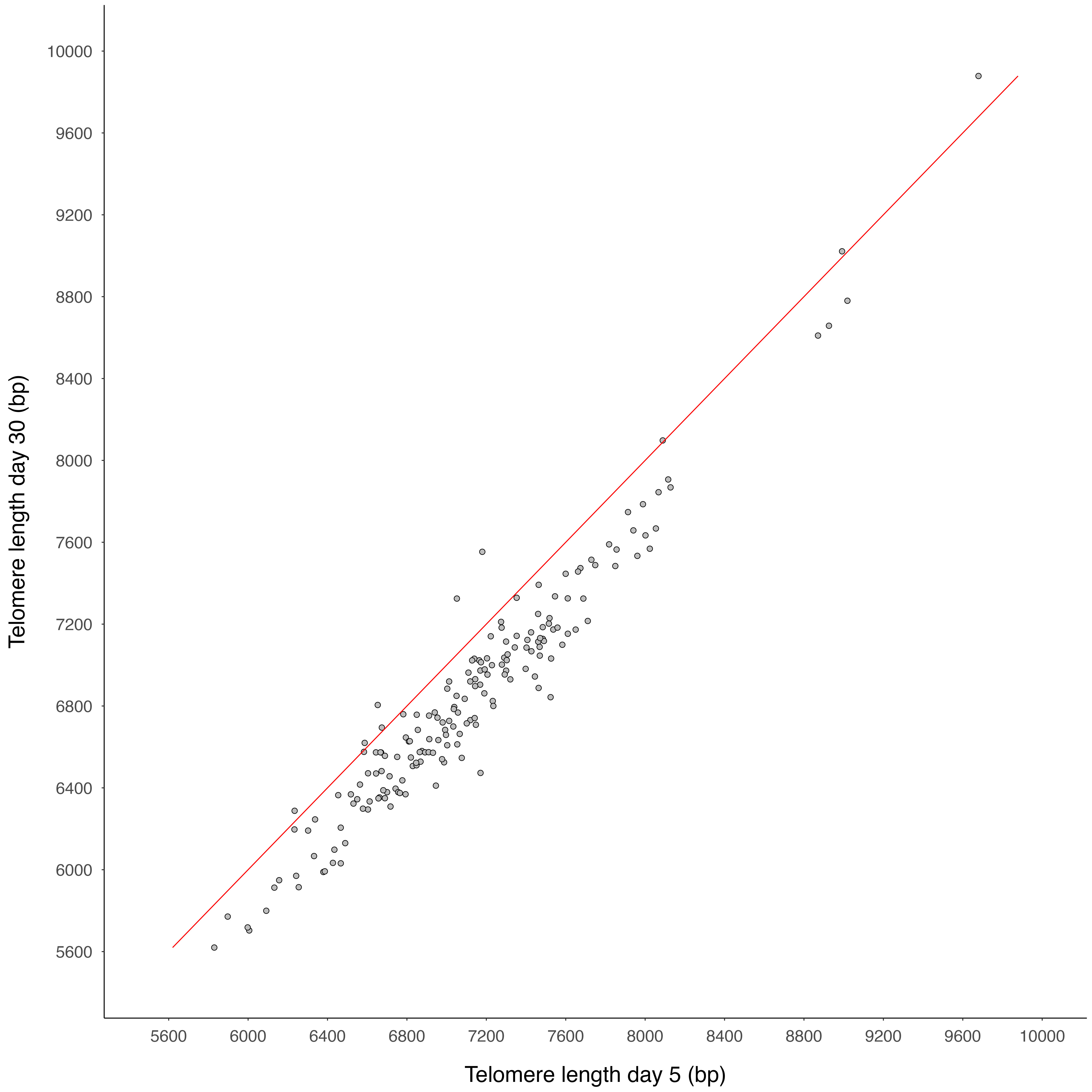
